# Knock-In of a 25-Kilobase Pair BAC-Derived Donor Molecule by Traditional and CRISPR/*Cas9*-Stimulated Homologous Recombination

**DOI:** 10.1101/076612

**Authors:** Tiffany Leidy-Davis, Kai Cheng, Leslie O. Goodwin, Judith L. Morgan, Wen Chun Juan, Xavier Roca, Sin-Tiong Ong, David E. Bergstrom

## Abstract

Here, we describe an expansion of the DNA size limitations associated with CRISPR knock-in technology, more specifically, the physical extent to which mouse genomic DNA can be replaced with donor (in this case, human) DNA at an orthologous locus. Driving our efforts was the desire to create a whole animal model that would replace 17 kbp of the mouse *Bcl2l11* gene with the corresponding 25-kbp segment of human *BCL2L11*, including a conditionally removable segment (2.9-kbp) of intron 2, a cryptic human exon immediately 3′ of this, and a native human exon some 20 kbp downstream. Using two methods, we first carried out the replacement by employing a combination of bacterial artificial chromosome recombineering, classic ES cell targeting, dual selection, and recombinase-driven cassette removal (traditional approach). Using a unique second method, we employed the same vector (devoid of its selectable marker cassettes), microinjecting it along with CRISPR RNA guides and *Cas9* into mouse zygotes (CRISPR approach). In both instances we were able to achieve humanization of *Bcl2l11* to the extent designed, remove all selection cassettes, and demonstrate the functionality of the conditionally removable, *loxP*-flanked, 2.9-kbp intronic segment.

**AUTHOR SUMMARY:** Clustered regularly interspaced short palindromic repeat (CRISPR) technology can be used to place DNA sequences (designed in the laboratory) into the genomes of living organisms. Here, we describe a new method, whereby we have replaced an exceptionally large segment of the mouse *Bcl2l11* gene with the corresponding segment of human *BCL2L11* gene. The method represents an expansion of the DNA size limitations typically associated with the introduction of DNA sequences through traditional CRISPR methods.

## INTRODUCTION

The discovery of clustered regularly interspaced short palindromic repeat (CRISPR) systems, the elucidation of their function, and their exploitation as genome engineering tools are revolutionizing genetic engineering (1–5). Discovered as a form of adaptive immunity in bacteria and archaea, CRISPR systems consist of a series of DNA spacer elements derived from invading plasmids or viruses. Interdigitated among the spacers is a series of direct repeats. Depending on the particular system, these series are transcribed and processed into single spacer/repeat units called crRNAs (CRISPR RNAs). In turn, these crRNAs may interact with other short RNAs (*e.g.*, tracrRNA) and one or more CRISPR-associated (Cas) proteins (*e.g.*, Cas9 of *Streptococcus pyogenes*), culminating in the assembly of an RNA-guided endonuclease directed at degrading DNA from the offending plasmid or virus.

As genome engineering tools, the CRISPR-Cas endonucleases serve as instruments for generating DNA double-strand breaks (DSBs) with locus-of-interest specificity, at high frequency, and across a wide variety of strains and organisms (6). When faced with DSBs, cells of the organism being perturbed respond with one or both of two DNA repair pathways known as the non-homologous end joining (NHEJ) pathway and the homology-directed repair (HDR) pathway (7, 8). DNA DSBs repaired by the more rapid and error-prone NHEJ pathway are characterized by the deletion or insertion of a small number of nucleotides. As one might expect, these insertion/deletion events (INDELS) within the open reading frame of a protein of interest may lead to the deletion of one or more endogenous amino acids, the insertion of one or more non-native amino acids, premature termination, or frameshift mutations. In each of these instances the modified mutant locus will commonly encode a hypomorphic or null allele of the original gene of interest.

In contrast, DSBs repaired in the presence of a homologous template (*e.g.*, sister chromatid, donor molecule) may be repaired by HDR (9). For genetic engineers, this provides the opportunity to introduce precise DNA modifications, created at the laboratory bench, into the organism under investigation, at the site of the DSB.

For traditional gene-targeting, of the sort in use in the mouse for the past thirty years (10–14), the traditional paradigm based on a large body of literature, has been to create plasmid vectors with two homology arms of a few to several kilobase pairs in length to act as donor molecules (15, 16). These arms are situated within the plasmid so as to flank investigator-altered sequences that will be incorporated into the genome after introduction of the plasmid vector into embryonic stem (ES) cells and HDR. Positive and negative selection cassettes are frequently employed to aid in selecting the rare ES cell clones containing properly integrated sequences. This technique is sufficient for modifying genomic sequence on a scale from one nucleotide to several thousand base pairs. The method may fall short, however, when attempting to alter entire mouse genes that often extend over 10s or 100s of thousands of base pairs.

In these instances, other genetic engineering technologies are employed including such methods as random transgenesis (17, 18), targeted transgenesis (19, 20), and recombinase-mediated cassette exchange (RMCE) (21, 22). Each of these methods has its drawbacks as well. For example, random transgenic methods deviate from genome modification at the cognate endogenous locus, sufficing to allow transgenes to integrate randomly (where they are subject to variegated expression). During targeted transgenesis, transgenes may be directed specifically to standardized safe harbor sites to limit this position-effect variegation but even here the transgenes are unlinked to their endogenous cognate genes. Like the related RMCE method, targeted transgenesis may involve the use of antibiotic selection cassettes flanked by recombinase-binding sites. In addition to the added complexity, deleting these selection cassettes requires breeding to specific recombinase-expressing mice thereby prolonging strain development (23–26).

With the advent of CRISPR technologies many new avenues have opened. For example, by dramatically increasing the frequency of DSBs at specified sites, gene-targeting need no longer be married to the culture of ES cells or the use and removal of selection cassettes. In fact, in mice, most experiments begin with the microinjection of *Cas9* and CRISPR RNA guides, (and when needed, donor molecules) into single-celled zygotes (27). Furthermore, in species where ES cell technology is lacking, CRISPR technology is a viable alternative, a fact that has opened gene-editing experimentation to a wide variety of strains and a broad range of species from bacteria to humans (6). However, here again, DNA modifications have generally been limited to physical extents on the order of a few to a few thousand base pairs. Moreover, systematic studies of the effect of homology arm length on CRISPR-associated HDR are lacking.

Despite the associated uncertainties, as described in the report that follows, we sought to expand the limits of CRISPR knock-in technology. Specifically, we attempted to increase the physical extent to which mouse genomic DNA could be replaced with donor (in this case, human) DNA at an orthologous locus. Driving our efforts was the desire to create a whole animal model that would replace 17 kbp of the mouse *Bcl2l11* gene with the corresponding segment of human *BCL2L11*, including a conditionally removable segment (2.9-kbp) of intron 2, a cryptic human exon immediately 3′ of this, and a native human exon some 20 kbp downstream (28). Using two approaches, we first carried out the replacement by employing a combination of bacterial artificial chromosome (BAC) recombineering, classic ES cell targeting, dual selection, and recombinase-driven cassette removal (hereafter referred to as our traditional approach) (29). In the second approach, we used the same vector (devoid of its selectable marker cassettes), microinjecting it along with CRISPR RNA guides and *Cas9* into mouse zygotes (hereafter referred to as our CRISPR approach). In both instances we were able to achieve humanization of *Bcl2l11* to the extent designed, remove all selection cassettes, and demonstrate the functionality of the conditionally removable, *loxP*-flanked, 2.9-kbp intronic segment.

Our latter result may well represent the largest segment of mouse DNA to be replaced by an orthologous human DNA using a CRISPR-directed approach with zygotic injection, to date. The study offers a proof-of-principal demonstration that a minimum of at least 25 kbp of genomic DNA can be effectively humanized in mouse, and provides a foundation for further technical optimization in mouse and specialization for use in other species.

## METHODS

### Husbandry

All mice were obtained from The Jackson Laboratory (Bar Harbor, ME), housed on a bedding of white pine shavings, and fed NIH-31 5K52 (6% fat) diet and acidified water (pH 2.5 to 3.0), *ad libitum*. All experiments were performed with the approval of The Jackson Laboratory Institutional Animal Care and Use Committee (IACUC) and in compliance with the Guide for the Care and Use of Laboratory Animals (8^th^ edition) and all applicable laws and regulations.

### Preparation of the Targeting Vectors/Donor Molecules

We designed a targeting vector/donor molecule with three objectives in mind — 1), to humanize a central segment of the *BCL2L11*/*Bcl2l11* gene; 2), to place selectable markers immediately 5’ and 3’ of the humanized segment; and 3), to flank a 2,903-bp region within one of the humanized introns with *loxP* sites in order to model a disease-associated deletion observed in 12% of the East Asian population (28).

Specifically, we constructed a targeting vector/donor molecule containing a 27,282-bp central segment of the human *BCL2L11* gene flanked by 12,773- and 26,632-bp homology arms consisting of the proximal and distal regions of the mouse *Bcl2l11* gene, respectively. This construct was designed such that it could be used both for homologous recombination in embryonic stem (ES) cells, as well as for a CRISPR/*Cas9* knock-in approach (See Fig 1).

**Fig 1.**
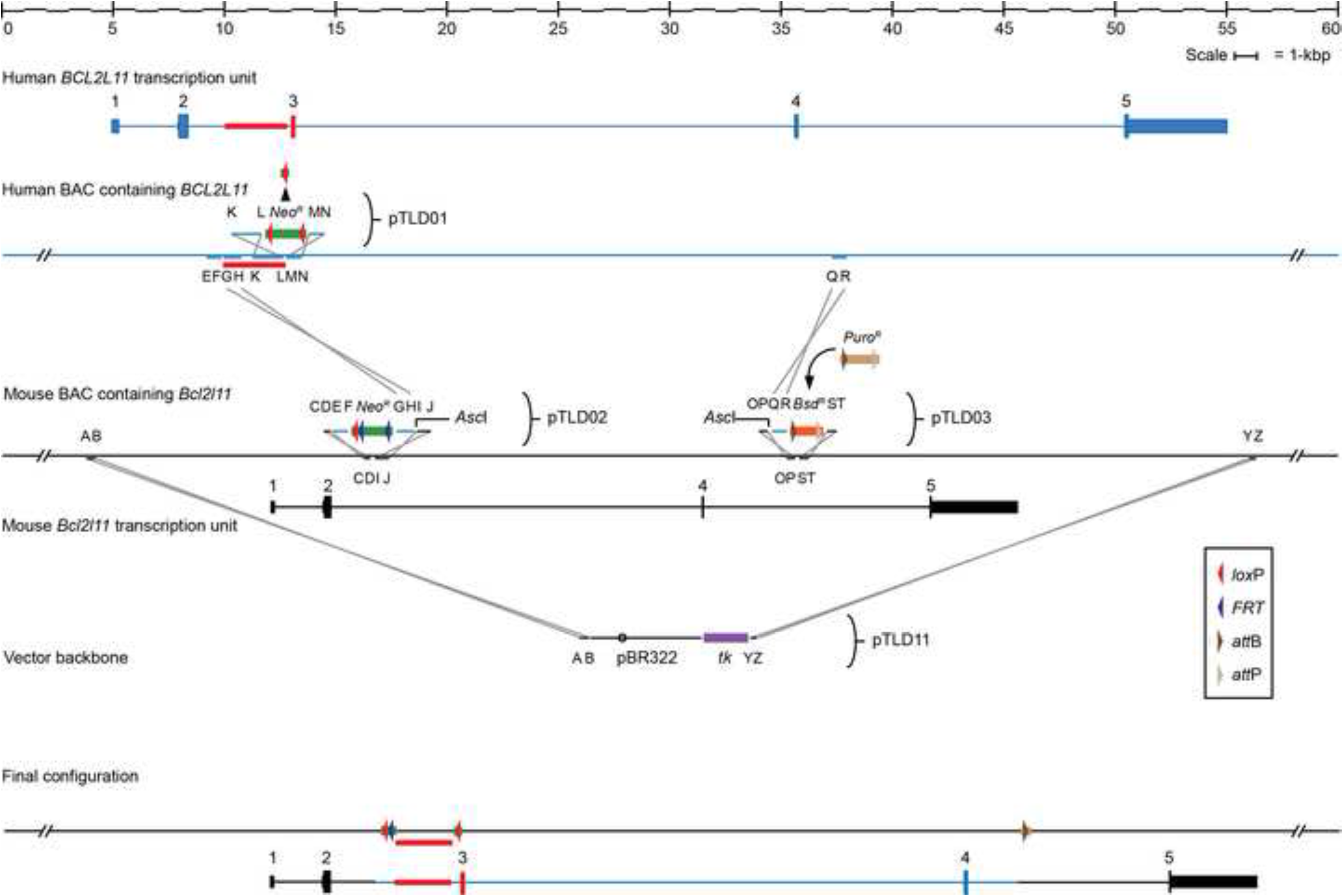
Construction of the *Bcl2l11/BCL2L11* targeting vector/donor molecule. A gene-targeting vector/donor molecule was constructed placing a 25-kbp segment of the human *BCL2L11* gene between mouse homology arms, placing removable selectable marker cassettes at each end of the human segment, and placing *loxP* sites around a 2.9-kbp segment of human DNA deleted in 12% of the East Asian population. See text for details.

Initially, BAC DNAs were purified from BAC clones containing the corresponding *BCL2L11* and *Bcl2l11* genes (human: library RP11, clone 695-B-23; mouse: library RP23, clone 331-K-22) (30, 31). Purified DNAs were then electroporated into the recombinogenic *E. coli* strain, SW102 (32).

Segments from the mouse and human BACs were amplified using the oligonucleotides described in Table 1, restriction-digested at sites incorporated into the oligonucleotides, gel-purified, and assembled into small plasmid vectors as follows:

**Table 1.** Oligonucleotides.

| Abbreviation | Synonym | Orientation | Species | Genome Build | Chromosomal Coordinates | Product Size | Homology Length | Overall Length | Sequence | Enzyme |
| --- | --- | --- | --- | --- | --- | --- | --- | --- | --- | --- |
| A | oTLD38 | + | Mouse | mm10 | Chr 2 : 128116776 128116795 | 424 | 20 | 31 | 5' -dCGCATACTAGTTCCATCCGGTCAATTCTCTC-3' | *SpeI* |
| B | oTLD39 | - | Mouse | mm10 | Chr 2 : 128117158 128117177 |  | 20 | 31 | 5' -dCGCATAAGCTTTTGTGTTGCTCCAGATTCC-3' | *HinDIII* |
| C | oTLD29new | + | Mouse | mm10 | Chr 2 : 128129327 128129348 | 246 | 22 | 35 | 5' -dCGCATGCGGCCGCATAGTTAATAACCACCAAGGCA-3' | *NotI* |
| D | oTLD30 | - | Mouse | mm10 | Chr 2 : 128129528 128129548 |  | 21 | 32 | 5' -dCGCATAAGCTTAAGTACTGACTGTAGCCCAAGAA-3' | *HinDIII* |
| E | oTLD27 | + | Human | hg38 | Chr 2 : 111124898 111124917 | 742 | 20 | 31 | 5' -dCGCATAAGCTTTATTGCTCAGAGGGTTTGA-3' | *HinDIII* |
| F | oTLD28 | - | Human | hg38 | Chr 2 : 111125598 111125617 |  | 20 | 31 | 5' -dCGCATGGATCCTGATTACCTCACTGAAGCC-3' | *BamHI* |
| G | oTLD20 | + | Human | hg38 | Chr 2 : 111125618 111125641 | 765 | 24 | 35 | 5' -dCGCATGAATTCGGCAGGCCCTTGCCCATGTTATAG-3' | *EcoRI* |
| H | oTLD21new | - | Human | hg38 | Chr 2 : 111126335 111126358 |  | 24 | 37 | 5' -dCGCATGGCGCCCTACTTTACTTCACAGGTATAACC-3' | *AscI* |
| I | oTLD31 | + | Mouse | mm10 | Chr 2 : 128129549 128129573 | 843 | 25 | 38 | 5' -dCGCATGGCGCCGCTAGAAATTTCTAAAACTATATTC-3' | *AscI* |
| J | oTLD32 | - | Mouse | mm10 | Chr 2 : 128130340 128130367 |  | 28 | 39 | 5' -dCGCATGTCGACGTATTAAGACTCTAATAGTCTCCAGAGG-3' | *SalI* |
| K | oTLD9B | + | Human | hg38 | Chr 2 : 111127109 111127130 | 1441 | 22 | 35 | 5' -dCGCATGCGGCCGCTCTCCTTACACTCTGGGAGGAT-3' | *NotI* |
| L | oTLD10 | - | Human | hg38 | Chr 2 : 111128507 111128525 |  | 19 | 30 | 5' -dCGCATGGATCCAACAGCATGATGGTTCCCC-3' | *BamHI* |
| M | oTLD11 | + | Human | hg38 | Chr 2 : 111128526 111128545 | 501 | 20 | 31 | 5' -dCGCATGAATTCCTCCATAGAGGCTGTGCCAT-3' | *EcoRI* |
| N | oTLD12 | - | Human | hg38 | Chr 2 : 111128985 111129004 |  | 20 | 31 | 5' -dCGCATGTCGACTGAGTGGGAAGAGTCAAGCC-3' | *SalI* |
| O | oTLD13 | + | Mouse | mm10 | Chr 2 : 128147195 128147214 | 478 | 20 | 33 | 5' -dCGCATGCGGCCGCTAAGGACCTCTCCCATCC-3' | *NotI* |
| P | oTLD14 | - | Mouse | mm10 | Chr 2 : 128147618 128147646 |  | 20 | 33 | 5' -dCGCATGGCGCCGCCCAACAGGACAGCCAGCTAC-3' | *AscI* |
| Q | oTLD15 | + | Human | hg38 | Chr 2 : 111151266 111151285 | 842 | 20 | 33 | 5' -dCGCATGGCGCCGCTGACTGCTTCCGCTAAAGG-3' | *AscI* |
| R | oTLD16 | - | Human | hg38 | Chr 2 : 111152064 111152083 |  | 20 | 31 | 5' -dCGCATGAATTCCTCCCACTTTGATCCTGAA-3' | *EcoRI* |
| S | oTLD17 | + | Mouse | mm10 | Chr 2 : 128147689 128147710 | 510 | 22 | 33 | 5' -dCGCATGGATCCGCATCTTCAGAAAGCAGTGTGTT-3' | *BamHI* |
| T | oTLD18 | - | Mouse | mm10 | Chr 2 : 128148155 128148176 |  | 22 | 33 | 5' -dCGCATGTCGACTCCTCAGTCCATTATCAACAG-3' | *SalI* |
| Y | oTLD40 | + | Mouse | mm10 | Chr 2 : 128173955 128173974 | 388 | 20 | 31 | 5' -dCGCATAAGCTTATCAGGCCAGGGTTCTAGT-3' | *HinDIII* |
| Z | oTLD41 | - | Mouse | mm10 | Chr 2 : 128174299 128174318 |  | 20 | 33 | 5' -dCGCATGCGGCCGCATAGTGTCCCTGTCCCAAGG-3' | *NotI* |

Oligonucleotides used in the construction of donor vectors. Restriction enzyme sites have been incorporated within 11- to 13-base segments of non-homology at the 5' end of each primer.

Segments KL and MN were cloned along with the neomycin resistance gene- (*Neo^R^*-) containing *Eco*RI/*Bam*HI fragment of PL452, into a pBluescript II vector (Agilent Technologies, Santa Clara, CA USA) modified to contain an R6Kγ origin of replication (33, 34). This plasmid is named pTLD01.

Segments CD, EF, GH, and IJ were cloned along with the neomycin resistance gene- (*Neo^R^*-) containing *Eco*RI/*Bam*HI fragment of PL451, into a pBluescript II (Agilent Technologies, Santa Clara, CA USA) vector modified to contain an R6Kγ origin of replication (33, 34). This plasmid is named pTLD02.

Segments OP, QR, and ST were cloned along with the blasticidin resistance gene- (*Bsd^R^*-) containing *Eco*RI/*Bam*HI fragment of pTLD08 (a PL452 derivative carrying *att*B, *att*P, and *Bsd^R^*), into a pBluescript II vector (Agilent Technologies, Santa Clara, CA USA) modified to contain an R6Kγ origin of replication (33, 34). This plasmid is named pTLD03.

Segments AB and YZ were cloned into a pBR322-based vector along with the negatively selectable thymidine kinase (*tk*) gene (35, 36). This plasmid is named pTLD11.

To begin the assembly of our humanized donor vector proper, pTLD01 was used with standard recombineering approaches to place a *loxP*-flanked neomycin resistance cassette (*Neo^R^*) just distal to the 2,903-bp deletion region in the human *BCL2L11*-containing BAC (29). After transferring the modified BAC to the *Cre*-expressing *E. coli* strain, SW106, the *Neo* cassette was removed by exposing cells to arabinose, leaving a single *loxP* site remaining (32).

Next, plasmid pTLD02 was used with standard recombineering techniques to place the EF segment of human DNA, a *loxP* site, an FRT-flanked *Neo* cassette, and the GH segment of human DNA just distal to mouse Exon 2 in the mouse *Bcl2l11*-containing BAC.

Next, plasmid pTLD03 was used with standard recombineering techniques to place the QR segment of human DNA, and an *attB/attP*-flanked blasticidin resistance (*Bsd^R^*) cassette, slightly distal to mouse Exon 4 in the pTLD02-modified, mouse *Bcl2l11*-containing BAC described above.

Next, plasmid pTLD11 was linearized with *Hind*III and used with standard recombineering procedures to retrieve the AB to YZ segment of the mouse *Bcl2l11* gene from the pTLD02/pTLD03-modified BAC, becoming pTLD14.

At this point, plasmid pTLD14 was purified, digested with *Asc*I, and its two major fragments resolved by agarose gel electrophoresis. The larger of the two linear fragments was gel-purified and electroporated into recombinogenic *E. coli* cells containing the *loxP*-modified human BAC clone described above, thus capturing the 27,282-bp human segment between flanking mouse homology arms, becoming plasmid pTLD15.

After experiencing some difficulty with blasticidin-based embryonic stem (ES) cell selection, we replaced the open reading frame (ORF) of *Bsd^R^* with that of *Puro*^*R*^ through a negatively selectable *rpsL* intermediate (37).

This completed vector, pTLD39, performed well in embryonic stem cells subjected to sequential neomycin/puromycin selection.

For the purpose of CRISPR/*Cas9*-based zygotic microinjection, the final *Neo^R^*/*Bsd^R^*-containing vector (plasmid pTLD15) was electroporated; first, into the FLP-expressing *E. coli* strain SW105 to remove *Neo^R^* (making plasmid pTLD66), and next, into a ϕC31 recombinase-expressing *E. coli* strain (an SW105 derivative) to remove *Bsd^R^* (32). The final vector was named pTLD67.

### Electroporation

For our traditional approach, we electroporated 25 µg of linear pTLD39 DNA into 1.5 × 10^7^ cells of the JM8-A3 (Strain: C57BL/6N) line of mouse embryonic stems cells. ES cells were then plated in ES+2i medium with sequential gentamycin (G418, 200 µg/ml, Gibco, Fisher Thermo Scientific, Waltham, MA, USA) and puromycin (0.75 µg/ml, Sigma-Aldrich, St. Louis, MO, USA) selection (38). Surviving ES cell clones were propagated on ES+2i medium, karyotyped, further tested for the presence of the puromycin resistance cassette by PCR, and assessed for homology arm, insert, and neomycin resistance cassette count by quantitative PCR. Properly targeted clones were microinjected into 3.5-days *post coitum* (dpc) blastocysts (see below).

### CRISPR sgRNA Design and Production

For our CRISPR approach, all single-guide RNAs (sgRNAs) were designed using an algorithm available at http://crispr.mit.edu (39). These sgRNAs, shown in Table 2, were designed along two concepts. In the first, the two highest scoring sgRNAs (one in each direction) within a 250-bp region were selected from both the 5’ and 3’ ends of the 17-kbp segment of the mouse *Bcl2l11* segment being replaced. In the second, two internal sgRNAs (one in each direction) closest to each end of the replaced segment were selected regardless of their overall score. Guides were produced according to the method of Briner, *et al*. (40). *Cas9* mRNA (CRISPR associated protein 9 mRNA, 5-methylcytidine, pseudouridine) was purchased from TriLink Biotechnologies (San Diego, CA).

**Table 2.**
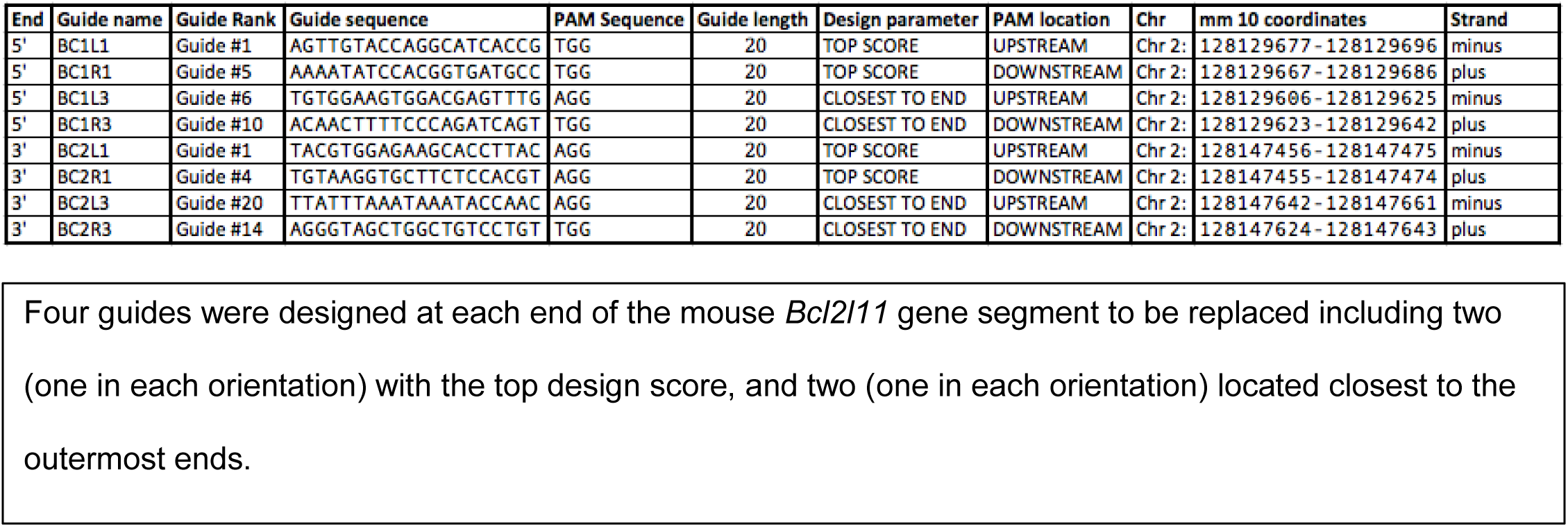
Single Guide RNAs (sgRNAs).

### Microinjection

For our traditional approach, properly targeted ES clones were microinjected into 3.5-dpc blastocysts, and the blastocysts transferred to pseudopregnant host dams, by standard techniques (41). The resulting embryos were allowed to go to term; the pups were delivered naturally and reared by the dams until weaning at four weeks of age.

For our CRISPR approach, microinjection mixes were prepared as shown in Table 3. Approximately 80 C57BL/6NJ zygotes were microinjected (in one to two technical replicates with each microinjection mix described above), transferred to pseudopregnant females by standard techniques, and allowed to go to term where they were reared by the dams until weaning at four weeks of age.

**Table 3.** Microinjection Mixes.

| Experiment | Number | 1 | 2 | 3 | 4 | 5 | 6 | 7 | 8 |
| --- | --- | --- | --- | --- | --- | --- | --- | --- | --- |
| Guides | BC1L1 | | | 50 ng/μL | 50 ng/μL | | | 50 ng/μL | 50 ng/μL |
| | BC1R1 | | | 50 ng/μL | 50 ng/μL | | | 50 ng/μL | 50 ng/μL |
| | BC1L3 | 50 ng/μL | 50 ng/μL | | | 50 ng/μL | 50 ng/μL | | |
| | BC1R3 | 50 ng/μL | 50 ng/μL | | | 50 ng/μL | 50 ng/μL | | |
| | BC2L1 | | | 50 ng/μL | 50 ng/μL | | | 50 ng/μL | 50 ng/μL |
| | BC2R1 | | | 50 ng/μL | 50 ng/μL | | | 50 ng/μL | 50 ng/μL |
| | BC2L3 | 50 ng/μL | 50 ng/μL | | | 50 ng/μL | 50 ng/μL | | |
| | BC2R3 | 50 ng/μL | 50 ng/μL | | | 50 ng/μL | 50 ng/μL | | |
| Other | Cas9 mRNA | 100 ng/μL | 100 ng/μL | 100 ng/μL | 100 ng/μL | 100 ng/μL | 100 ng/μL | 100 ng/μL | 100 ng/μL |
| Reagents | Donor vector | 5 ng/μL | 1 ng/μL | 5 ng/μL | 1 ng/μL | 10 ng/μL | 5 ng/μL | 10 ng/μL | 5 ng/μL |
| Embryos | Rep 1 | ~80 | ~80 | ~80 | ~80 | ~80 |  | ~80 |  |
| Injected | Rep 2 |  |  |  |  |  | ~80 |  | ~80 |

Microinjection mixes contained four guides (either those with the highest scores or those with the most terminal positions within the mouse Bcl2l11 segment to be replaced) and varying concentrations of donor DNA (1, 5, or 10 ng/ μL).

### Genotyping

Potentially chimeric mice, arising from the microinjection of 3.5-dpc blastocysts (traditional approach) or 1-celled zygotes (CRISPR approach), and their progeny were genotyped at designed Cas9 binding sites using the oligonucleotide primers described in Table 4. As shown (See Fig 2), these primers were used in pairs, in separate PCR reactions designed to amplify DNA across: 1) the Cas9 binding sites of intact (or small INDEL-containing) mouse alleles, 2) the mouse/human junctions of humanized alleles (or randomly integrating transgenes), and 3) the breakpoints of any deletion-bearing alleles.

**Table 4.** Genotyping oligonucleotides.

| Primer Name | Sequence | Length (nt) | Forward/Reverse | Chromosome | Coordinates (mm10/hg38) |
| --- | --- | --- | --- | --- | --- |
| oTLD56 | 5' -dATCTGTGGCCTTCTAGCCAA-3' | 20 | Forward | Mouse Chr 2 | 128129240-128129259 |
| oTLD57 | 5' -dAGAATGCCCTAACTCAGCCA-3' | 20 | Reverse | Mouse Chr 2 | 128130445-128130464 |
| oTLD239 | 5' -dTGCATCTAAGGGTTTGGCTT-3' | 20 | Forward | Mouse Chr 2 | 128147294-128147313 |
| oTLD338 | 5' -dGAGTCAAAGCCTACATCCCCAA-3' | 22 | Reverse | Mouse Chr 2 | 128147780-128147801 |
| oTLD335 | 5' -dGGAACAGCAAGTCGATCAACAC-3' | 22 | Reverse | Human Chr 2 | 111125334-111125355 |
| oTLD337 | 5' -dGGTGTTTGAGGAGAGTGCTGTA-3' | 22 | Forward | Human Chr 2 | 111151728-111151749 |

Standard PCR primers were designed to amplify the junctions flanking the original mouse *Bcl2l11* allele, the humanized *BCL2L11* allele, and the deletion-bearing allele.

**Fig 2.**
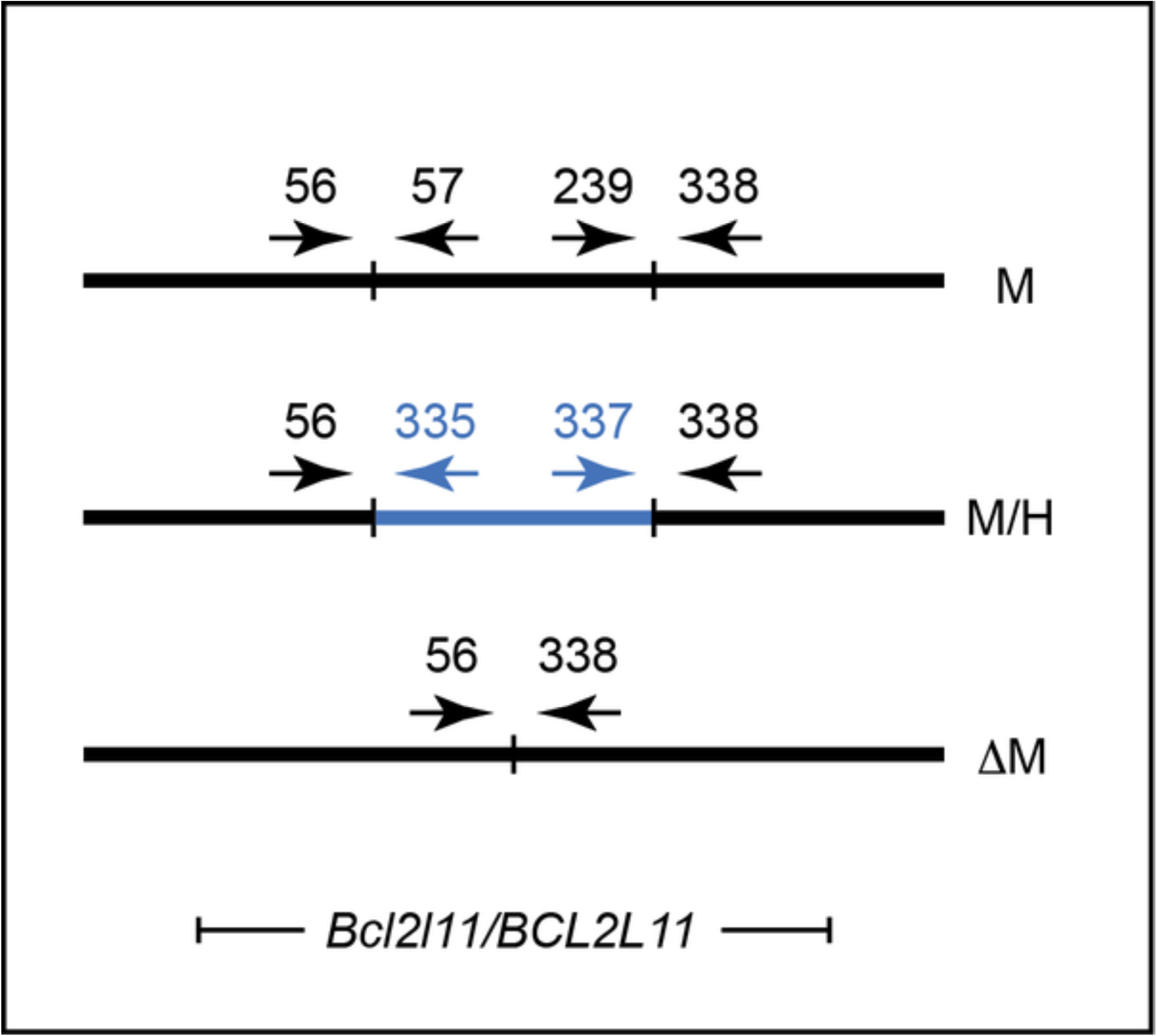
Organization of genotyping primers for mouse (M), humanized (M/H), and deletion-bearing (M) alleles of *BCL2L11/Bcl2l11*. Schematic showing the organization of genotyping primers. Numbers, primer designation as in Table 4; left and right segments of horizontal black lines, flanking regions of the mouse *Bcl2l11* region; central segment of top horizontal black line, central (to be replaced) region of the mouse *Bcl2l11* gene; blue line, central segment of human *BCL2L11* gene. See text for details.

### Sanger Sequencing

For more detailed analysis of specific alleles, PCR products from genotyping reactions were purified and sequenced by JAX Scientific Services according to the method developed by Sanger (42). PCR products were purified using HighPrep PCR magnetic beads (MagBio Genomics, Gaithersburg, MD USA). Cycle sequencing was performed using a BigDye Terminator Cycle Sequencing Kit, version 3.1 (Applied Biosystems, Foster City, CA USA). Sequencing reactions contained 5µl of purified PCR product (3-20 ng) and 1µl of primer at a concentration of 5 pmol/µl. Sequencing reaction products were purified using HighPrep DTR (MagBio Genomics, Gaithersburg, MD USA). Purified reactions were run on an Applied Biosystems 3730xl DNA Analyzer (Applied Biosystems, Foster City, CA USA). Sequence data were analyzed using Sequencing Analysis Software, version 5.2 (Applied Biosystems, Foster City, CA USA). Resulting sequence (.abi) files were imported into Sequencher, version 5.0.1 (Gene Codes Corporation, Ann Arbor, MI USA), for further analysis.

### Genetic mapping

To show that the human segment of *BCL2L11* had replaced its mouse counterpart in the orthologous *Bcl2l11* locus, we used genetic mapping to localize the humanized segment of the *BCL2L11/Bcl2l11* gene (See Fig 3). Two backcrosses were established using the following approach. First, FVB/NJ females were crossed to C57BL/6NJ males carrying the humanized segment to obtain F_1_ hybrid (FVBB6NF1/J) progeny. These progeny were then genotyped for the presence of the humanized segment. Males carrying the human sequence (FVBB6NF1/J-*BCL2L11*) were backcrossed to either FVB/NJ females or C57BL/6NJ females to generate N_2_ progeny.

**Fig 3.**
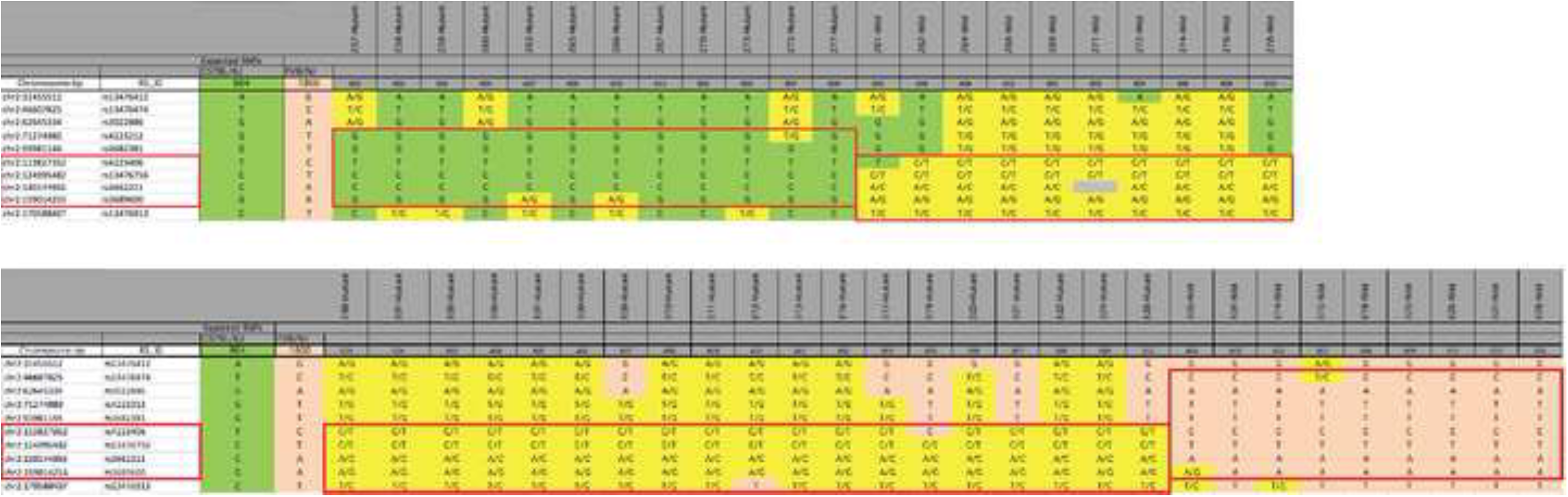
Linkage analysis of the *BCL2L11* integration site following CRISPR-stimulated homologous recombination in mouse zygotes. Shown are the linkage analyses for 22 F2 progeny of a C57BL/6NJ X FVBB6NF1/J-*BCL2L11* backcross (upper panel) and 28 F2 progeny of an FVB/NJ X FVBB6NF1/J-*BCL2L11* backcross (lower panel). Linkage and haplotype analysis indicate that the *BCL2L11* vector’s integration has occurred between markers rs4223406 and rs3689600 and its segregation is fully concordant with markers rs13476756 and rs3662211. This result is entirely consistent with integration of the human *BCL2L11* segment within the endogenous mouse *Bcl2l11* gene as designed.

These backcross schemes can be annotated as follows:

C57BL/6NJ X FVBB6NF1/J-*BCL2L11*
FVB/NJ X FVBB6NF1/J-*BCL2L11*

N_2_ progeny from each backcross (along with appropriate controls) were genotyped using KASP-chemistry (LGC Limited, Teddington, UK) across a set of approximately 150 single-nucleotide polymorphism (SNP) markers distributed roughly equally across the mouse genome. Concordance between each marker in the set and the humanized segment was calculated by chi-square (χ^□^) analysis.

## RESULTS

### Traditional Approach

Following electroporation of the pTLD39 vector into the JM8-A3 line of ES cells and selection on G418, we assayed 89 surviving clones for the presence of the puromycin resistance cassette by PCR. Of these, twenty-seven contained the puromycin cassette and were subjected to puromycin selection. Of these, four clones survived and were assessed for homology arm, insert, and neomycin resistance cassette count by quantitative PCR. One clone passed all of these tests for proper targeting of the central human *BCL2L11* segment to the endogenous mouse *Bcl2l11* gene. ES cells from this clone were microinjected into blastocysts resulting in nine high-quality chimeras. The four highest quality male chimeras were mated to C57BL/6NJ females resulting in two independent instances of germline transmission of the humanized allele. Although presumably identical, independent lines (genetic background: C57BL/6JN) were developed from each instance. Mating males with B6N.Cg-Tg(*Sox2-Cre*)1Amc/J female mice resulted in progeny in which the *loxP*-flanked 2.9-kbp human intronic segment was deleted, as designed.

### CRISPR Approach

At term, a total of 94 pups were born; six were stillborn and six did not survive to four weeks of age. Eighty-two mice were weaned and distributed among experiments as shown in Table 4.

Both Experiment 3 (highest scoring guides, 5 ng/µL donor DNA) and Experiment 5 (guides closest to ends, 10 ng/µL donor DNA) resulted in no viable pups remaining at wean-age. Despite these results Experiment 7 (conducted with a donor DNA concentration equal to that of Experiment 5, *i.e.*, 10 ng/µL) and Experiment 8 (a replicate of Experiment 3) resulted in seven and 21 pups, respectively, suggesting that the lack of pups in Experiments 3 and 5 was due to technical failure rather than anything systematically wrong with the experimental design.

To genotype these 82 progeny PCR assays were designed to span each of the proximal and distal mouse/human junctions and to span the 17-kbp mouse region to be replaced. The results of these experiments are shown in Table 5. As noted, PCR assays designed to span each of the proximal and distal mouse/human junctions identified three founders that were positive for both (Experiment 2, guides closest to ends, 1 ng/µL donor DNA; Experiment 6, guides closest to ends, 5 ng/µL donor DNA; and Experiment 7, highest scoring guides, 10 ng/µL donor DNA). PCR assays designed to span the 17-kbp mouse region to be replaced identified two of the three founders described above (Experiment 6, guides closest to ends, 5 ng/µL donor DNA; and Experiment 7, highest scoring guides, 10 ng/µL donor DNA).

**Table 5.**
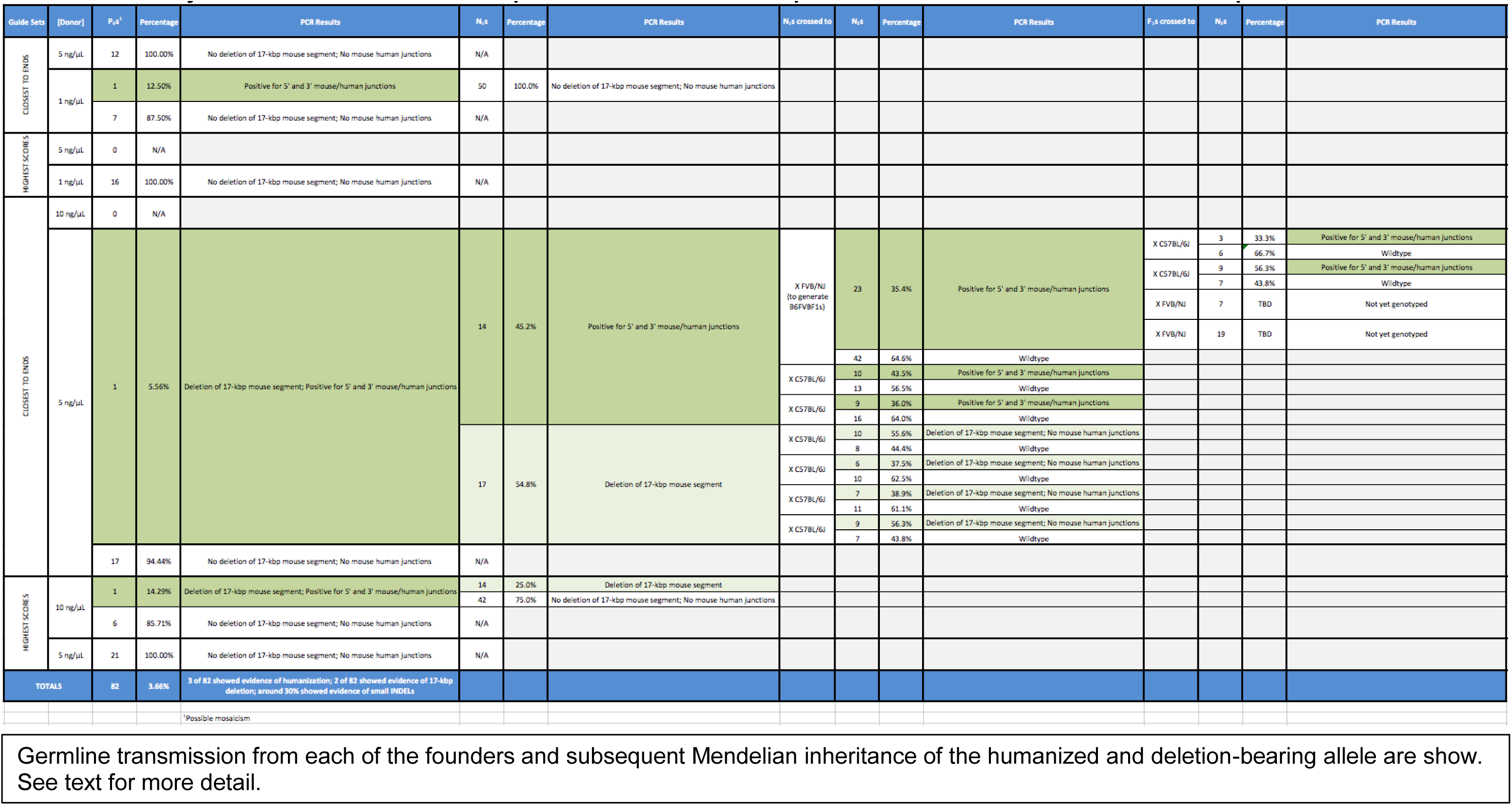
Summary of the CRISPR-stimulated replacement of 17-kilobase pairs of mouse *Bcl2l11* with 25-kilobase pairs of human *BCL2l11*.

To further explore the inheritance of these genetic changes, we mated the human insertion/deletion-positive P_0_s from Experiments 2, 6, and 7 to C57BL/6J mice and genotyped their progeny. The results of these analyses are shown in Table 5. As shown, the human insertion-positive P_0_ mouse (male) from Experiment 2 (guides closest to ends, 1 ng/µL donor DNA) failed to transmit the humanized allele to any of 29 of its N_1_ progeny suggesting that the P_0_ mouse is mosaic with a germline consisting primarily of unmodified wildtype cells.

In contrast, the human insertion- and deletion-positive P_0_ mouse (male) from Experiment 7 (highest scoring guides, 10 ng/µL donor DNA) transmitted its deletion-bearing allele to four of its 21 N_1_ progeny. This P_0_ mouse, however, did not transmit the human insertion-bearing allele to any of these 21 mice again suggesting that the P_0_mouse is mosaic with a germline consisting of relatively few human insertion-bearing cells.

Interestingly, the human insertion- and deletion-positive P_0_ mouse (female) from Experiment 6 (guides closest to ends, 5 ng/µL donor DNA) transmitted either a human insertion-bearing allele or a deletion-bearing allele to all of its 13 N_1_ progeny, but never both, implying that this animal is breeding as a true heterozygote with a genotype of both human insertion- and deletion-bearing alleles at the *Bcl2l11* locus. Subsequent breeding of three select N_1_ mice (two bearing the human insertion and one bearing the deletion) gave results consistent with Mendelian expectations. Mating males with B6N.Cg-Tg(*Sox2-Cre*)1Amc/J female mice resulted in progeny in which the *loxP*-flanked 2.9-kbp human intronic segment was deleted, as designed.

### Genetic mapping

We used an outcross-backcross genetic mapping strategy as a means of localizing the insertion site of BAC-derived human *BCL2L11* sequences. Twenty-two N_2_ progeny were analyzed from the C57BL/6NJ X (FVB/NJ X C57BL/6NJ) backcross and twenty-eight progeny from the FVB/NJ X (FVB/NJ X C57BL/6NJ) backcross. Analysis of the data demonstrates strong linkage between the human *BCL2L11* segment and several genetic markers on mouse Chromosome 2 (See Fig 3). In the backcross to C57BL/6NJ, the marker with strongest linkage, marker rs13476756, had a log-odds ratio (LOD) of 6.58 (p<0.004). In the backcross to FVB/NJ, marker rs13476756 had a LOD score of 7.64 (p<0.0004).

Analysis of individual haplotypes (specifically, points of recombination in samples 261, 263, 266, 303, and 319) further narrows the insertion-critical region to a 45.2-Mbp region from marker rs4223406 (nucleotide 113,827,352) to marker rs3689600 (nucleotide 159,014,253) on Mouse Chromosome 2 (GRC38/mm10), which is consistent with integration into the 36,510-bp mouse *Bcl2l11* gene that spans from nucleotide 128,126,038 to nucleotide 128,162,547. Put another way, this analysis shows that both the mouse *Bcl2l11* gene and the engineered human sequences must be colocalized within a region comprising less than 2% of the mouse genome. We conclude that integration of the human sequence has not occurred randomly, but has indeed occurred by homologous recombination as designed.

## DISCUSSION

Contemporary CRISPR technology is revolutionizing genetic engineering and has contributed [along with zinc-finger nuclease (ZFN) and transcription activator-like effector nuclease (TALEN) technologies] to the newly emergent field of gene editing (43–56). The greater CRISPR technique is in a period of rapid expansion, its methodology now being applied across dozens of species in thousands of laboratories around the globe.

Moreover, the seminal core technology continues to diversify with additional enzymatic reagents, novel applications, and technical improvements under robust investigation. This, in turn, has led to a rapid expansion of CRISPR knowledge and the publication of CRISPR reports and reviews on a daily basis.

In the experiments reported here, we set out to explore the feasibility of using CRISPR technology to replace large (10s of kbp) segments of the mouse genome with human DNA from orthologous loci. Current CRISPR approaches aimed at knocking experimental DNAs into a locus of interest by homologous recombination have generally involved relatively small genomic expanses from single nucleotides to a few kilobase pairs. Moreover, these experiments routinely make use of long oligonucleotides, or targeting vectors with sub-kilobase homology arms, as donor molecules. Only more rarely are targeting vectors used of the sizes routinely employed in studies involving mouse ES cells.

In contrast to these common practices, we surmised that experimentally altered DNAs, of 10s to 100s of kilobase pair lengths, might be directed into a locus of interest if the DNA were outfitted with homology arms 15-30 times longer than those in common use today. Accordingly, we used a traditional approach with bacterial artificial chromosomes containing both human and mouse genomic DNA to prepare a donor molecule with 25-kbp of human *BCL2L11* genomic sequence flanked by 15-kbp and 30-kbp mouse homology arms. We have demonstrated by PCR, sequence, and linkage analysis that replacement of a minimum of 25-kbp of mouse genomic DNA can be achieved using human DNA from the corresponding locus.

Given this proof-of-principle, future studies can now begin to explore questions of efficiency and optimization. In our experiment we performed microinjection into mouse zygotes to see if mice could be recovered, with any degree of humanization of the mouse *Bcl2l11* gene, and if these mice were capable of transmitting the humanized allele through the germline to their offspring. These experiments have demonstrated the feasibility of this CRISPR/BAC technology to introduce experimental DNA in a directed fashion to the zygotic genome and the ability of the specifically targeted DNA to be transmitted through the germline to progeny. However, due to the small number of data points in whole animal experiments, one can only speculate on the impact of guide selection and donor DNA concentration variables on overall success rates.

Among the experiments in which donor DNA was detected in P_0_ mice (Experiments 2, 6, and 7), DNA donor concentrations of 1, 5, and 10 ng/µL were represented but the resulting mice show varying degrees of mosaicism. In Experiment 2, where donor DNA concentration was at its lowest (1 ng/µL), donor DNA was not detected among N_1_ progeny (0/50) suggesting that integration of the donor DNA occurred at multicellular stage of embryonic development and that those cells that did acquire the donor DNA did not contribute to the germline at an appreciable level.

In Experiment 6, where donor DNA concentration was at an intermediate level (5 ng/µL), donor DNA was detected among nearly half of all N_1_ progeny (14/31) suggesting that integration of the donor DNA occurred at the one-cell (zygotic) stage of embryonic development, that that cell gave rise to all cells of the germline, and that the donor DNA was passed, during meiosis, into half of the population of mature spermatozoa. This result is consistent with our hypothesis that a deletion, of the 17-kbp mouse segment to be replaced, occurred at the *Bcl2l11* locus in the homologous chromosome in the zygote, and was transmitted, in repulsion to the DNA insertion, to all remaining progeny (17/17). This result is entirely congruent with the optimal desired outcome, *i.e.*, where the P_0_ zygote undergoes biallelic modification, develops into a mouse with no mosaicism, and transmits one or the other variant alleles in equal numbers (50%:50%) to the population of mature spermatozoa.

In Experiment 7, where donor DNA concentration was at the highest level tested (10 ng/µL), the 17-kbp deletion was detected in only 25% of all N_1_ progeny (14/56), and the donor DNA, present in the P_0_ mouse, was not transmitted to the N1 generation at all (0/56). These results can be explained assuming a scenario whereby a deletion occurred in one *Bcl2l11* allele, in a single blastomere, at or near the two-cell stage, and that this deletion-bearing cell gave rise to roughly half of the developing premeiotic germline and a fourth of all mature (postmeiotic) germcells. At some later point in blastogenesis, one can hypothesize that an insertion of donor DNA occurred, but in so few cells as to not contribute to the germline in an appreciable way.

A number of aspects in Experiment 7 may have contributed to its less than optimal result. First, due to its viscosity, a donor DNA preparation with a DNA concentration that is too high may not be efficiently delivered through the microinjection needle to the zygote, or delivered in a form less conducive to promoting Cas9 activity and/or HDR. Moreover, the guides designed for this experiment, although designed to have an optimal score, did not have what we surmised to be an optimal position, near the ends of the mouse DNA segment to be replaced. It may be that, in experiments of this type, guide position represents a more significant design parameter than guide score alone. It is interesting to note that, among all experiments using guides designed for high score optimization, only in Experiment 7, where donor DNA concentration was at the highest level tested (10 ng/µL), was any evidence of donor DNA incorporation seen, and even here it was at a level apparently so low in the P_0_ founder mouse as to not transmit the modified allele to N_1_ mice. You may recall that, in the previously mentioned Experiment 6, where an optimal result *was* achieved, donor DNA concentration was only 5 ng/µL. It is entirely possible that the successful result seen in that instance was driven by superiorly performing/positioned (nearest the end) guides even at what could prove to be a suboptimal donor DNA concentration. Comparing Experiment 6 with Experiment 7, it is interesting to note that the experiment with the higher donor DNA concentration (Experiment 7, 10 ng/µL) did achieve a higher rate of incorporation (as a percentage of live born mice, 14.3% versus 5.6%) but a lower quality of allele modification in the single founder recovered (mosaicism/transmission of only one modified allele at low frequency compared to nonmosaicism/transmission of both modified alleles at maximum frequency). One may speculate that DNA concentration may be the most important parameter related to the introduction of DNA into individual zygotes; whereas, guide design may prove to be the most important factor for promoting more frequent deletion formation and more efficient HDR once donor DNA has entered the cell. Further experimentation, performed in large numbers of cells *in vitro*, is likely to be a productive avenue for optimization of this technique.

## ACKNOWLEDGMENTS

The authors thank the staff of The Jackson Laboratory’s Scientific Research Services in both the Genetic Engineering Technologies and Reproductive Sciences groups. The authors thank Dr. Narayanan Raghupathy, The Jackson Laboratory, for assistance with statistical analyses. The authors thank Jennifer Cook and Louise Dionne for mouse colony management as well as Rachel Urban, Susan Kales, and Jiayuan Shi for their molecular biological expertise.

## REFERENCES

1. Wright AV, Nunez JK, Doudna JA. Biology and Applications of CRISPR Systems: Harnessing Nature’s Toolbox for Genome Engineering. Cell. 2016 Jan 14;164(1-2):29–44. PubMed PMID: 26771484.

2. Sternberg SH, Doudna JA. Expanding the Biologist’s Toolkit with CRISPR-Cas9. Molecular cell. 2015 May 21;58(4):568–74. PubMed PMID: 26000842.

3. Jiang F, Doudna JA. The structural biology of CRISPR-Cas systems. Current opinion in structural biology. 2015 Feb;30:100–11. PubMed PMID: 25723899. Pubmed Central PMCID: 4417044.

4. Hochstrasser ML, Doudna JA. Cutting it close: CRISPR-associated endoribonuclease structure and function. Trends in biochemical sciences. 2015 Jan;40(1):58–66. PubMed PMID: 25468820.

5. Doudna JA, Charpentier E. Genome editing. The new frontier of genome engineering with CRISPR-Cas9. Science. 2014 Nov 28;346(6213):1258096. PubMed PMID: 25430774.

6. Sander JD, Joung JK. CRISPR-Cas systems for editing, regulating and targeting genomes. Nature biotechnology. 2014 Apr;32(4):347–55. PubMed PMID: 24584096. Pubmed Central PMCID: 4022601.

7. Davis AJ, Chen DJ. DNA double strand break repair via non-homologous end-joining. Translational cancer research. 2013 Jun;2(3):130–43. PubMed PMID: 24000320. Pubmed Central PMCID: 3758668.

8. Morrical SW. DNA-pairing and annealing processes in homologous recombination and homology-directed repair. Cold Spring Harbor perspectives in biology. 2015 Feb;7(2):a016444. PubMed PMID: 25646379.

9. Lundgren M, Charpentier E, Fineran PC, editors. CRISPR: Methods and Protocols: Springer; 2015.

10. Capecchi MR. Altering the genome by homologous recombination. Science. 1989 Jun 16;244(4910):1288–92. PubMed PMID: 2660260.

11. Capecchi MR. The new mouse genetics: altering the genome by gene targeting. Trends in genetics : TIG. 1989 Mar;5(3):70–6. PubMed PMID: 2660363.

12. Capecchi MR. Targeted gene replacement. Scientific American. 1994 Mar;270(3):52–9. PubMed PMID: 8134827.

13. Capecchi MR. Choose your target. Nature genetics. 2000 Oct;26(2):159–61. PubMed PMID: 11017070.

14. Capecchi MR. Gene targeting in mice: functional analysis of the mammalian genome for the twenty-first century. Nature reviews Genetics. 2005 Jun;6(6):507–12. PubMed PMID: 15931173.

15. Bouabe H, Okkenhaug K. A protocol for construction of gene targeting vectors and generation of homologous recombinant embryonic stem cells. Methods in molecular biology. 2013;1064:337–54. PubMed PMID: 23996269. Pubmed Central PMCID: 4526796.

16. Bouabe H, Okkenhaug K. Gene targeting in mice: a review. Methods in molecular biology. 2013;1064:315–36. PubMed PMID: 23996268. Pubmed Central PMCID: 4524968.

17. Palmiter RD, Brinster RL. Germ-line transformation of mice. Annual review of genetics. 1986;20:465–99. PubMed PMID: 3545063.

18. Palmiter RD, Brinster RL. Transgenic mice. Cell. 1985 Jun;41(2):343–5. PubMed PMID: 2985274.

19. Ohtsuka M. Development of pronuclear injection-based targeted transgenesis in mice through Cre-loxP site-specific recombination. Methods in molecular biology. 2014;1194:3–19. PubMed PMID: 25064095.

20. Ohtsuka M, Ogiwara S, Miura H, Mizutani A, Warita T, Sato M, et al. Pronuclear injection-based mouse targeted transgenesis for reproducible and highly efficient transgene expression. Nucleic acids research. 2010 Dec;38(22):e198. PubMed PMID: 20880997. Pubmed Central PMCID: 3001095.

21. Turan S, Galla M, Ernst E, Qiao J, Voelkel C, Schiedlmeier B, et al. Recombinase-mediated cassette exchange (RMCE): traditional concepts and current challenges. Journal of molecular biology. 2011 Mar 25;407(2):193–221. PubMed PMID: 21241707.

22. Turan S, Zehe C, Kuehle J, Qiao J, Bode J. Recombinase-mediated cassette exchange (RMCE) - a rapidly-expanding toolbox for targeted genomic modifications. Gene. 2013 Feb 15;515(1):1–27. PubMed PMID: 23201421.

23. Hayashi S, Lewis P, Pevny L, McMahon AP. Efficient gene modulation in mouse epiblast using a Sox2Cre transgenic mouse strain. Mechanisms of development. 2002 Dec;119 Suppl 1:S97–S101. PubMed PMID: 14516668.

24. Hayashi S, Tenzen T, McMahon AP. Maternal inheritance of Cre activity in a Sox2Cre deleter strain. Genesis. 2003 Oct;37(2):51–3. PubMed PMID: 14595839.

25. Rodriguez CI, Buchholz F, Galloway J, Sequerra R, Kasper J, Ayala R, et al. High-efficiency deleter mice show that FLPe is an alternative to Cre-loxP. Nature genetics. 2000 Jun;25(2):139–40. PubMed PMID: 10835623.

26. Raymond CS, Soriano P. High-efficiency FLP and PhiC31 site-specific recombination in mammalian cells. PloS one. 2007;2(1):e162. PubMed PMID: 17225864. Pubmed Central PMCID: 1764711.

27. Qin W, Kutny PM, Maser RS, Dion SL, Lamont JD, Zhang Y, et al. Generating Mouse Models Using CRISPR-Cas9-Mediated Genome Editing. Current protocols in mouse biology. 2016;6(1):39–66. PubMed PMID: 26928663. Pubmed Central PMCID: 4848752.

28. Ng KP, Hillmer AM, Chuah CT, Juan WC, Ko TK, Teo AS, et al. A common BIM deletion polymorphism mediates intrinsic resistance and inferior responses to tyrosine kinase inhibitors in cancer. Nature medicine. 2012 Apr;18(4):521–8. PubMed PMID: 22426421.

29. Copeland NG, Jenkins NA, Court DL. Recombineering: a powerful new tool for mouse functional genomics. Nature reviews Genetics. 2001 Oct;2(10):769–79. PubMed PMID: 11584293.

30. Osoegawa K, Tateno M, Woon PY, Frengen E, Mammoser AG, Catanese JJ, et al. Bacterial artificial chromosome libraries for mouse sequencing and functional analysis. Genome research. 2000 Jan;10(1):116–28. PubMed PMID: 10645956. Pubmed Central PMCID: 310499.

31. Osoegawa K, Mammoser AG, Wu C, Frengen E, Zeng C, Catanese JJ, et al. A bacterial artificial chromosome library for sequencing the complete human genome. Genome research. 2001 Mar;11(3):483–96. PubMed PMID: 11230172. Pubmed Central PMCID: 311044.

32. Warming S, Costantino N, Court DL, Jenkins NA, Copeland NG. Simple and highly efficient BAC recombineering using galK selection. Nucleic acids research. 2005;33(4):e36. PubMed PMID: 15731329. Pubmed Central PMCID: 549575.

33. Kolter R, Inuzuka M, Helinski DR. Trans-complementation-dependent replication of a low molecular weight origin fragment from plasmid R6K. Cell. 1978 Dec;15(4):1199–208. PubMed PMID: 728998.

34. Liu P, Jenkins NA, Copeland NG. A highly efficient recombineering-based method for generating conditional knockout mutations. Genome research. 2003 Mar;13(3):476–84. PubMed PMID: 12618378. Pubmed Central PMCID: 430283.

35. Balbas P, Soberon X, Bolivar F, Rodriguez RL. The plasmid, pBR322. Biotechnology. 1988;10:5–41. PubMed PMID: 3061523.

36. Balbas P, Soberon X, Merino E, Zurita M, Lomeli H, Valle F, et al. Plasmid vector pBR322 and its special-purpose derivatives–a review. Gene. 1986;50(1-3):3–40. PubMed PMID: 3034735.

37. Wang S, Zhao Y, Leiby M, Zhu J. A new positive/negative selection scheme for precise BAC recombineering. Molecular biotechnology. 2009 May;42(1):110–6. PubMed PMID: 19160076. Pubmed Central PMCID: 2669495.

38. Czechanski A, Byers C, Greenstein I, Schrode N, Donahue LR, Hadjantonakis AK, et al. Derivation and characterization of mouse embryonic stem cells from permissive and nonpermissive strains. Nature protocols. 2014 Mar;9(3):559–74. PubMed PMID: 24504480. Pubmed Central PMCID: 4112089.

39. Zhang F. CRISPR Design: Massachusettes Institute of Technology; 2015 [cited 2016 07/01/2016]. Available from: http://crispr.mit.edu.

40. Briner AE, Donohoue PD, Gomaa AA, Selle K, Slorach EM, Nye CH, et al. Guide RNA functional modules direct Cas9 activity and orthogonality. Molecular cell. 2014 Oct 23;56(2):333–9. PubMed PMID: 25373540.

41. Behringer R, Gertsenstein M, Vintersten-Nagy K, Nagy A. Manipulating the Mouse Embryo: A Laboratory Manual. 4th Edition ed. Cold Spring Harbor, NY: Cold Spring Harbor Laboratory Press; 2014.

42. Sanger F, Nicklen S, Coulson AR. DNA sequencing with chain-terminating inhibitors. Proceedings of the National Academy of Sciences of the United States of America. 1977 Dec;74(12):5463–7. PubMed PMID: 271968. Pubmed Central PMCID: 431765.

43. Heidenreich M, Zhang F. Applications of CRISPR-Cas systems in neuroscience. Nature reviews Neuroscience. 2016 Jan;17(1):36–44. PubMed PMID: 26656253. Pubmed Central PMCID: 4899966.

44. Hsu PD, Lander ES, Zhang F. Development and applications of CRISPR-Cas9 for genome engineering. Cell. 2014 Jun 5;157(6):1262–78. PubMed PMID: 24906146. Pubmed Central PMCID: 4343198.

45. Shalem O, Sanjana NE, Zhang F. High-throughput functional genomics using CRISPR-Cas9. Nature reviews Genetics. 2015 May;16(5):299–311. PubMed PMID: 25854182. Pubmed Central PMCID: 4503232.

46. Zhang F, Wen Y, Guo X. CRISPR/Cas9 for genome editing: progress, implications and challenges. Human molecular genetics. 2014 Sep 15;23(R1):R40-6. PubMed PMID: 24651067.

47. Gaj T, Gersbach CA, Barbas CF, 3rd. ZFN, TALEN, and CRISPR/Cas-based methods for genome engineering. Trends in biotechnology. 2013 Jul;31(7):397–405. PubMed PMID: 23664777. Pubmed Central PMCID: 3694601.

48. Ousterout DG, Gersbach CA. The Development of TALE Nucleases for Biotechnology. Methods in molecular biology. 2016;1338:27–42. PubMed PMID: 26443211.

49. Sommer D, Peters AE, Baumgart AK, Beyer M. TALEN-mediated genome engineering to generate targeted mice. Chromosome research : an international journal on the molecular, supramolecular and evolutionary aspects of chromosome biology. 2015 Feb;23(1):43–55. PubMed PMID: 25596827.

50. Sun N, Zhao H. Transcription activator-like effector nucleases (TALENs): a highly efficient and versatile tool for genome editing. Biotechnology and bioengineering. 2013 Jul;110(7):1811–21. PubMed PMID: 23508559.

51. Wright DA, Li T, Yang B, Spalding MH. TALEN-mediated genome editing: prospects and perspectives. The Biochemical journal. 2014 Aug 15;462(1):15–24. PubMed PMID: 25057889.

52. Carroll D. Genome engineering with zinc-finger nucleases. Genetics. 2011 Aug;188(4):773–82. PubMed PMID: 21828278. Pubmed Central PMCID: 3176093.

53. Durai S, Mani M, Kandavelou K, Wu J, Porteus MH, Chandrasegaran S. Zinc finger nucleases: custom designed molecular scissors for genome engineering of plant and mammalian cells. Nucleic acids research. 2005;33(18):5978–90. PubMed PMID: 16251401. Pubmed Central PMCID: 1270952.

54. Handel EM, Cathomen T. Zinc-finger nuclease based genome surgery: it’s all about specificity. Current gene therapy. 2011 Feb;11(1):28–37. PubMed PMID: 21182467.

55. Palpant NJ, Dudzinski D. Zinc finger nucleases: looking toward translation. Gene therapy. 2013 Feb;20(2):121–7. PubMed PMID: 22318089.

56. Swarthout JT, Raisinghani M, Cui X. Zinc Finger Nucleases: A new era for transgenic animals. Annals of neurosciences. 2011 Jan;18(1):25–8. PubMed PMID: 25205916. Pubmed Central PMCID: 4117018.

